# Two decades of EU pesticide substitution have not reduced impacts on stream invertebrates

**DOI:** 10.64898/2026.11.05.764581

**Authors:** Jonas Gröning, Matthias Liess

## Abstract

Chemical regulation assumes that withdrawing hazardous substances and replacing them with alternatives reduces environmental harm. We tested this premise within the EU pesticide regulatory framework using a 23-year record of event-driven pesticide exposure and stream-invertebrate communities from 152 German agricultural streams sampled between 1998 and 2021. Neither pesticide toxic pressure nor pesticide-related invertebrate impairment showed a detectable decline over time, and an improvement equivalent to one ecological status class can be excluded. Dominant toxicity drivers shifted from organophosphates to neonicotinoids and subsequently to pyrethroids, yet the relationship between toxic pressure and community effects remained consistent despite their distinct modes of action. Sustained toxic pressure reflected three mechanisms: successive chemical substitution, prolonged approval periods, and continued exposure to substances no longer authorised as plant protection products. Substitution thus changed which chemicals harm streams, not how much. Measured field outcomes must become the benchmark of pesticide regulation.

---

A central premise of chemical regulation is that withdrawing hazardous substances and replacing them with alternatives reduces harm to people and ecosystems. Yet replacements have repeatedly proved similarly problematic: bisphenol A was replaced by bisphenols S and F with comparable hormonal activity ^1^, long-chain per- and polyfluoroalkyl substances by structurally similar fluorinated alternatives ^2^, and polybrominated diphenyl ethers by organophosphate flame retardants ^3^. Evidence for this pattern, termed regrettable substitution ^4^, has relied almost entirely on comparisons of the hazard properties of substitutes. Whether successive substitutions ultimately reduce adverse effects has, to our knowledge, not been evaluated in the field.

Pesticides are a critical test case. They are designed to be biologically active, are released directly into the environment and are the dominant stressor for vulnerable invertebrates in agricultural streams ^5^. In the European Union, active substances are approved for up to ten years and renewed for up to fifteen years after reassessment against current scientific standards; if a renewal decision is delayed, the approval can be extended ^6^. Over the past three decades, EU pesticide regulation has restricted or withdrawn major insecticides, including the withdrawal of parathion in 2002 ^7^, the prohibition of all outdoor uses of clothianidin, imidacloprid and thiamethoxam in 2018 ^8–10^, and the non-renewal of thiacloprid in 2020 ^11^, consistent with the expectation that approved alternatives are less harmful ^12^. Official indicators suggest substantial progress: the Harmonised Risk Indicator 1 (HRI1), which weights active-substance sales by regulatory category, fell from 100 in the 2011–2013 baseline to 41 in the EU and 55 in Germany in 2024 ^13^. This downward trend appears to align with the EU objective of reducing the use and risk of chemical pesticides by 50% by 2030 relative to a 2015– 2017 baseline ^14^.

However, whether these regulatory measures have actually reduced the environmental impact cannot be answered with routine monitoring data. Monitoring under the Water Framework Directive underrepresents small agricultural streams that are particularly exposed to pesticide inputs, relies largely on grab samples that miss short, runoff-driven concentration peaks, and covers only a fraction of currently authorised pesticides ^5,15^. Sales-based indicators, in turn, cannot directly capture ecological effects in aquatic ecosystems, where exposure is shaped by episodic inputs and environmental fate, and community responses by substance-specific toxicity, multiple stressors and ecological interactions.

To provide field-based evidence, we compiled a 23-year record, to our knowledge unique in Europe, combining event-driven pesticide measurements with invertebrate community assessments from 152 German agricultural streams sampled in four campaigns between 1998 and 2021. Parts of these data have been published ^5,16–18^; here, we harmonised and reanalysed the data using a common set of effect concentrations, a common treatment of non-detects and a common effect indicator, and linked them to the authorisation histories of the substances. Because active-substance approval is harmonised at EU level, and because the indicator of pesticide effects used here, SPEAR_pesticides_, responds consistently to pesticide toxicity across Europe ^19^, this record serves as a sentinel for the EU authorisation system.

We ask (i) whether impairment of stream invertebrates and underlying pesticide toxic pressure declined over 23 years, (ii) whether exposure–effect relationships changed as dominant insecticides were replaced, and (iii) how shifts in toxicity drivers aligned with regulatory status.

## Results

### No detectable decline in invertebrate impairment or pesticide toxic pressure over 23 years

Pesticide-related impairment of invertebrate communities, quantified as normalised SPEAR_pesticides_ and adjusted for the proportion of cropland in the catchment, did not differ among the four campaigns (joint F-test, F = 1.19, df1 = 3, df2 = 95.0, p = 0.32; Figure 1a). Estimated marginal means ranged from 0.43 to 0.56 across campaigns, with values of 0.46 (95% CI: 0.36 to 0.57) in 1998–2000, 0.56 (0.47 to 0.65) in 2013, 0.53 (0.48 to 0.57) in 2018–2019 and 0.43 (0.29 to 0.57) in 2021. In every campaign, the average stream thus retained only about half of the proportion of pesticide-vulnerable taxa expected without pesticide exposure, corresponding to moderate status. The linear trend (+0.0004 units per year, 95% CI: −0.0059 to 0.0068; p = 0.89) corresponds to a change of +0.01 (−0.13 to +0.15) over the 22 years between the first and last campaign midpoints. An improvement by one ecological status class (0.2 units) therefore lies outside the confidence interval.

**Figure 1:**
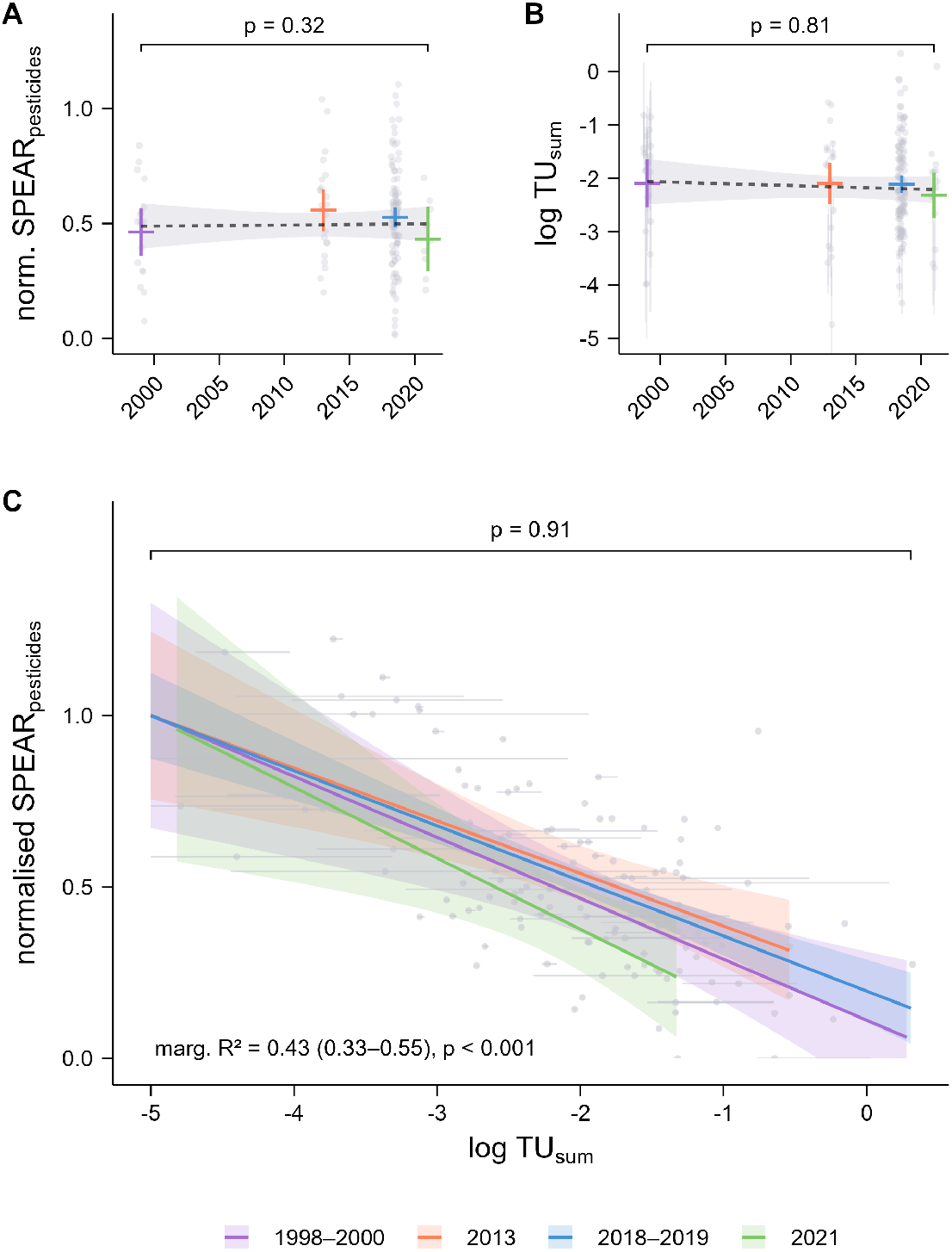
Pesticide-related invertebrate impairment and toxic pressure showed no detectable decline over 23 years, and the exposure–effect relationship remained consistent. a,b, Campaign-specific estimated marginal means and 95% confidence intervals of normalised SPEAR_pesticides_ (a) and log TU_sum_ (b); dashed lines and grey bands show the linear temporal trend and its 95% confidence interval; p values refer to the campaign effect. c, Campaign-specific exposure–effect relationships; coloured lines and bands indicate fixed-effect predictions and their 95% confidence intervals; the p value refers to the log TU_sum_ × campaign interaction. Grey points show site-level observations; grey lines indicate the 2.5th–97.5th percentile range of log TU_sum_ across imputations. SPEAR_pesticides_ was normalised to the campaign-specific prediction at log TU_sum_ = −5 (reference values: 0.33, 0.33, 0.35 and 0.28 for 1998– 2000, 2013, 2018–2019 and 2021, respectively). n = 18, 21, 101 and 10 sites in 1998–2000, 2013, 2018–2019 and 2021, respectively.

Toxic pressure, expressed as the logarithm of summed toxic units (log TU_sum_) and adjusted for the proportion of cropland in the catchment, likewise did not differ among campaigns (pooled Wald test, F = 0.33, df1 = 3, df2 = 167.1, p = 0.81; Figure 1b). Estimated marginal means ranged from −2.3 to −2.1 across campaigns, with values of −2.1 (95% CI: −2.5 to −1.6) in 1998–2000, −2.1 (−2.5 to −1.7) in 2013, −2.1 (−2.3 to −2.0) in 2018–2019 and −2.3 (−2.8 to −1.9) in 2021. The linear trend of −0.007 log units per year (95% CI: −0.031 to 0.018; p = 0.58) corresponds to a change in log TU_sum_ of −0.15 (−0.68 to +0.39) over 22 years, that is, a change in toxic pressure by a factor of 0.7 (95% CI: a 4.8-fold decrease to a 2.5-fold increase). Because community effects scale with the logarithm of toxic pressure, such changes are best judged by their ecological consequence: according to the field exposure–effect relationship, the estimated trend in log TU_sum_ corresponds to an increase in SPEAR_pesticides_ of 0.03 (95% CI: −0.07 to +0.12). Even the largest decrease compatible with the data would have improved SPEAR_pesticides_ by at most about 0.12, less than one ecological status class.

Results were similar for the subset of 21 sites sampled in multiple campaigns, thereby reducing differences in site composition among campaigns (SPEAR_pesticides_: +0.002 units per year, 95% CI: −0.005 to 0.009, p = 0.56; log TU_sum_: −0.005 log units per year, 95% CI: −0.038 to 0.028, p = 0.76; Supplementary Figure 2).

### Consistent exposure–effect relationships despite substitution

SPEAR_pesticides_ declined with increasing log TU_sum_ (marginal R2 = 0.43, 95% CI: 0.33–0.55; p < 0.001; Figure 1c). Campaign-specific slopes ranged from −0.21 to −0.15 per log unit (1998–2000: −0.18, −0.28 to −0.08; 2013: −0.15, −0.23 to −0.07; 2018–2019: −0.16, −0.20 to −0.12; 2021: −0.21, −0.35 to −0.06) and did not differ among campaigns (pooled Wald test, F = 0.17, df1 = 3, df2 = 134.8, p = 0.91). Accordingly, the relationship between pesticide toxic pressure and pesticide-related community effects remained consistent across monitoring campaigns, showing that similar levels of toxic pressure were associated with similar effects throughout the monitoring period.

### Shifts in toxicity drivers following regulatory action

Insecticides accounted for 88–97% of toxic pressure in all campaigns (Figure 2). In 1998–2000, the organophosphate parathion (34% mean contribution to TU_sum_, 2.5th–97.5th percentile range: 17– 50%) was the dominant toxicity driver; pyrethroids accounted for about half of the estimated toxic pressure; however, their concentrations were entirely below the limit of detection and therefore estimated by imputation (Supplementary Table 5). After parathion authorisations were withdrawn in 2002, organophosphates accounted for at most 2% of total toxic pressure. Following the introduction of thiacloprid, thiamethoxam, clothianidin and acetamiprid in the early 2000s, neonicotinoids had become the dominant contributors to invertebrate toxic pressure by 2013 (66%, 58–74%; thiacloprid 44%, 37–52%; clothianidin 16%, 10–24%). This pattern persisted in 2018–2019, when neonicotinoids again represented the largest share (72%, 71–72%; thiacloprid 46%, 45–46%; clothianidin 13%, 12–13%). After the ban of the neonicotinoids clothianidin, imidacloprid and thiamethoxam for outdoor use at the end of 2018 and the non-renewal of thiacloprid in 2020, the contribution of neonicotinoids declined (22%, 20–24%) in 2021, whereas the share of pyrethroids increased (from 20% in 2013 to 50%, 48–53%; lambda-cyhalothrin 26%, 23–30%; esfenvalerate 10%, 10–11%).

**Figure 2:**
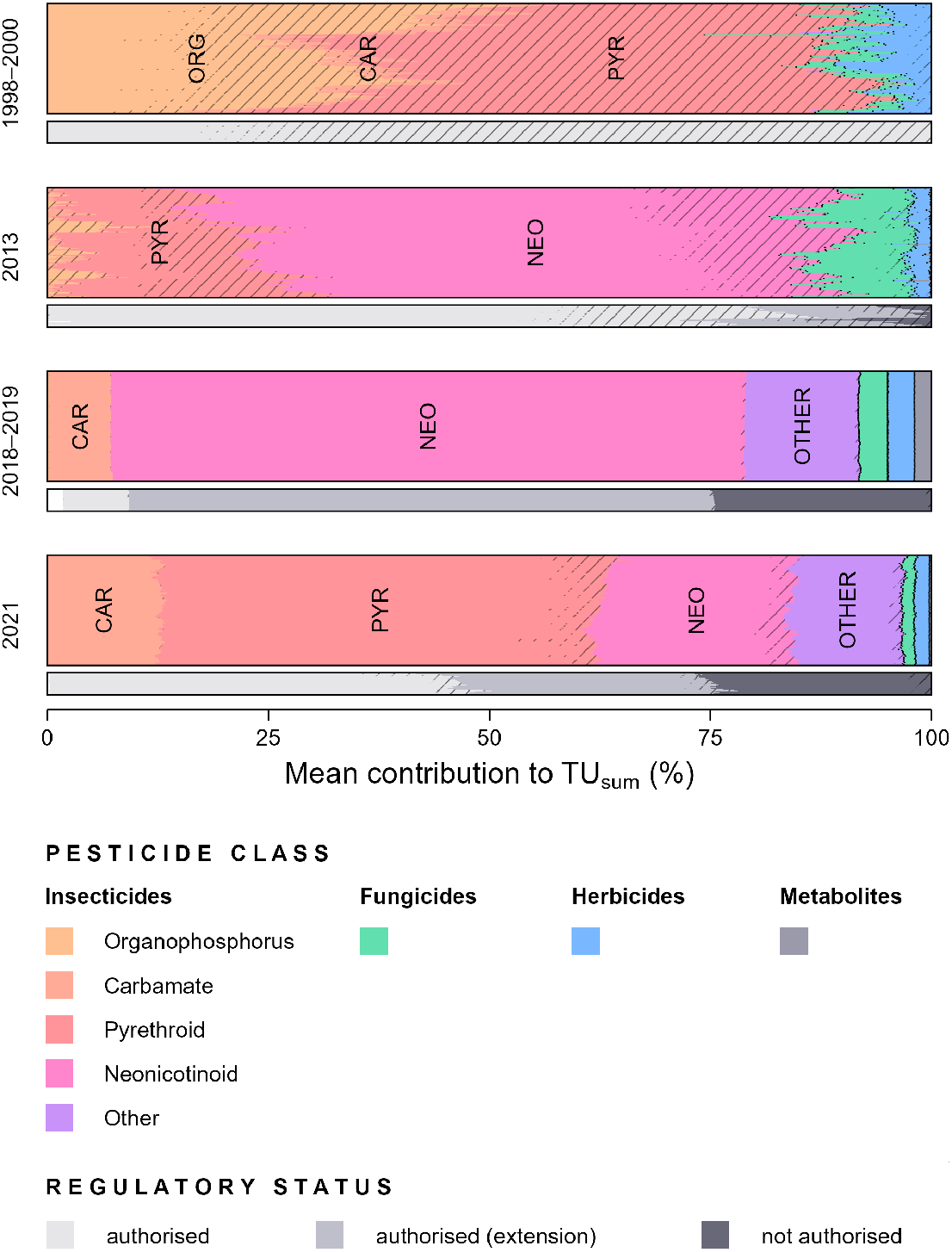
The substances driving invertebrate toxicity shifted, and a growing share came from substances outside regular approval. Mean relative contributions to TU_sum_ per campaign. Upper bars show contributions by pesticide type (insecticides, fungicides, herbicides, metabolites) and insecticide class (ORG, organophosphates; CAR, carbamates; PYR, pyrethroids; NEO, neonicotinoids); lower bars, by regulatory status at the time of sampling (authorised; authorised under extended approval; not authorised). Thin lines show individual imputations. Hatched areas indicate contributions from imputed concentrations. Only sites with ≥20% cropland in the catchment were included. n = 18, 17, 100 and 10 sites in 1998–2000, 2013, 2018–2019 and 2021, respectively.

The share of toxic pressure attributable to substances outside regular approval increased over time (Figure 2). In 1998–2000, all invertebrate toxic pressure stemmed from authorised substances, and in 2013 substances without authorisation contributed to a maximum of 4% (2.5th–97.5th percentile range: 0–13%). By 2018–2019, substances under extended approval or no longer authorised accounted for 91% (90–92%) of toxic pressure, and still for 54% (49–60%) in 2021; extended approvals contributed 66% (65–66%) and 29% (27–32%), respectively, while non-authorised substances contributed 25% (22–28%) in both campaigns. In 2018–2019, the extended-approval share was dominated by thiacloprid (45%, 44–45%) and clothianidin (9%, 8–9%), whose original approval periods had ended in 2014 and 2016, respectively. The non-authorised share was driven mainly by fipronil (13%), covered only by emergency authorisations from 2010 to 2015 ^20^, and by imidacloprid (7%, 7–8%) and clothianidin (4%), both no longer authorised after late 2018. In 2021, pirimicarb (13%, 12–13%), thiacloprid (9%, 8–10%) and cypermethrin (5%, 3–8%) contributed most under extended approval, four to five years after the end of their original approval periods, whereas fipronil (12%, 11–13%), permethrin (5%, 4–6%), clothianidin (4%, 3–5%) and imidacloprid (3%, 2– 3%) were the main non-authorised contributors.

### Prolonged approvals are the rule rather than the exception

Of the 113 measured active substances covered by Regulation (EC) No 1107/2009, 104 had their approval extended beyond the original expiry date (Figure 3). Cumulative extensions lasted 6.4 years on average; two-thirds exceeded five years and 9% exceeded ten years. Of the 59 extended substances for which a renewal decision has since been taken, 46% were finally not renewed.

**Figure 3:**
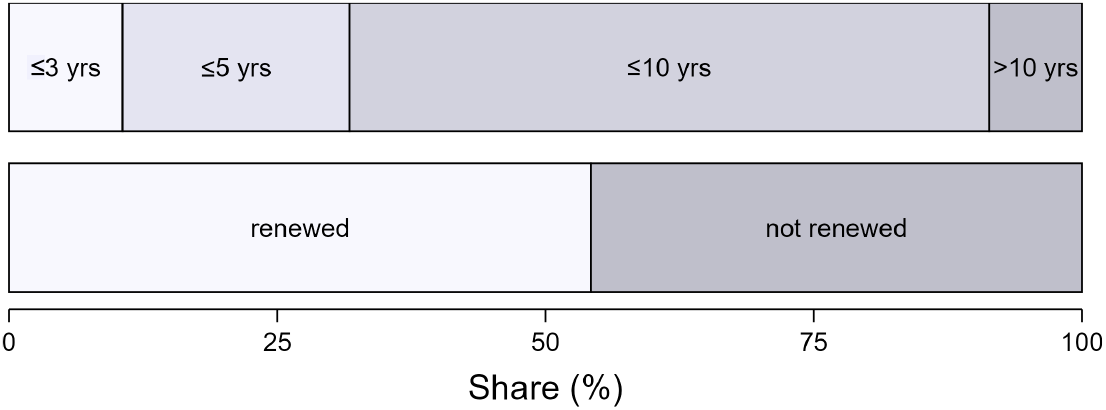
Approval extensions were widespread and frequently preceded non-renewal. Upper bar, distribution of cumulative approval-extension durations among the 104 measured active substances whose approval was extended beyond the original expiry date; lower bar, renewal outcomes for the 59 extended substances for which a renewal decision had been completed.

## Discussion

### Regulatory action resulted in substitution rather than risk reduction

Our results provide, to our knowledge, the first field evidence that pesticide substitutions following regulatory action have not reduced impacts on stream invertebrates. Earlier evidence of regrettable substitution rests on the hazard properties of substitutes ^1,3^, and analyses of pesticide use have shown that the toxicity applied through replacements can match or exceed that of banned compounds ^21,22^. Field studies have documented ecological damage after new insecticide classes were introduced ^23^, but not whether successive substitutions change the outcome over time. Here, these consequences were assessed directly in receiving ecosystems: across 23 years and successive shifts among three dominant insecticide classes, neither pesticide-related invertebrate impairment nor pesticide toxic pressure showed a detectable decline, and an improvement equivalent to even one ecological status class can be excluded.

Three mechanisms sustained toxic pressure. First, substitution: each withdrawal of a dominant insecticide was followed by the emergence of another, from parathion to neonicotinoids and, following neonicotinoid restrictions, to pyrethroids. Paralleling national sales ^20^, this sequence is consistent with substitution by other chemical insecticides rather than a transition to non-chemical pest control ^24^. Second, prolonged approvals: approval periods were extended for almost all measured active substances, by more than six years on average, while nearly half of those since reassessed were ultimately not renewed. Substances for which continued approval was no longer justified under current standards thus remained on the market, and in the streams, for years.

Recently, the General Court of the European Union clarified that such extensions under Article 17 of Regulation (EC) No 1107/2009 must remain temporary and exceptional (PAN Europe v Commission, T-412/22; Pollinis France v Commission, T-94/23). Third, persistence beyond authorisation: substances no longer authorised as plant protection products contributed a quarter of toxic pressure in 2018–2019 and 2021. Clothianidin and imidacloprid were still detected three years after their outdoor ban, consistent with their persistence in soils ^25^. Similar legacy effects have been reported from Danish agricultural streams, where legacy pesticides accounted for about 20% of summed toxic units in storm-flow samples ^26^.

### Consistent exposure–effect relationship despite changing toxicity drivers

Within the EU regulatory framework, periodic reassessment is intended to incorporate new scientific knowledge into approval decisions, while the European Commission has argued that newer active substances are expected to have improved toxicological and ecotoxicological properties ^12^. However, the substitution of dominant insecticidal substances neither reduced toxic pressure in the field (Figure 1b) nor altered the exposure–effect relationship (Figure 1c). This consistency is not self-evident, as the dominant toxicity drivers shifted from organophosphates to neonicotinoids and subsequently to pyrethroids, three classes with distinct modes of action: organophosphates inhibit acetylcholinesterase, neonicotinoids act on nicotinic acetylcholine receptors and pyrethroids modulate voltage-gated sodium channels ^27^. These mechanistic differences could be expected to produce different sensitivity profiles and, consequently, different community-level responses. However, the classes share an overlapping group of sensitive and pesticide-vulnerable insect taxa ^16,27^, providing a plausible ecological explanation for why comparable levels of toxic pressure were associated with similar community-level effects across monitoring campaigns. An insecticide that replaces a banned compound must control the same pests and is therefore likely to harm the same non-target insects: harm follows insecticidal function rather than chemical identity. Likewise, comparable exposure–effect relationships have been reported across Europe, Siberia and Australia ^19^, as well as in the Argentine Pampas ^28^ and East Africa ^29^, where SPEAR_pesticides_ required only minor adaptations.

Beyond this, the stability of the exposure–effect relationship shows that toxic units derived from standard acute tests remained a valid, substance-independent measure of ecological effects across insecticide classes. Moreover, the relationship confirms that community-level effects in the field occur at exposure levels considered acceptable in authorisation: at log TU_sum_ = −2, equivalent to 1/100 of the EC50 for a single substance, which corresponds to the first-tier acute regulatory acceptable concentration, the proportion of pesticide-vulnerable taxa is reduced to about 50% of reference conditions. This finding is supported by a joint analysis of field studies from three continents, which found effects on sensitive taxa at 1/1,000 to 1/10,000 of the EC50 of Daphnia magna ^19^, as well as by evidence that regulatory thresholds for invertebrate toxicity drivers may be insufficiently protective by a factor of 5.3 to 40 ^5^.

### Robustness of the findings

Although all monitoring campaigns targeted small streams in agricultural landscapes and combined event-driven pesticide sampling with macroinvertebrate community assessments, direct comparisons were constrained by differences in site composition, analyte coverage and generally higher detection limits in the earlier campaigns. We addressed differences in site composition by accounting for agricultural land-use intensity in comparisons among monitoring campaign means and in trend analyses, as pesticide toxic pressure increased with the proportion of cropland (Supplementary Figure 3). Restricting the analyses to the 21 sites sampled in more than one campaign, thereby reducing differences in site composition among campaigns, yielded the same overall conclusion of no temporal decline in pesticide-related invertebrate impairment or pesticide toxic pressure (Supplementary Figure 2). In each monitoring campaign, target substances were selected based on regional or national use or sales information and expected toxicity to aquatic invertebrates to capture substances of high environmental relevance. The main limitation in analyte coverage was the lack of pyrethroid measurements in 2018–2019 due to analytical constraints. This, however, can only have led to an underestimation of toxic pressure in that campaign, by a factor of about 1.2 if pyrethroids contributed as much as in 2013, and therefore cannot explain the absence of a temporal decline. Neonicotinoids were not analysed in 1998–2000; however, only imidacloprid was then authorised, and even with its 2013 share (5%, 2–10%), toxic pressure in 1998–2000 would have been underestimated by a factor of at most 1.1. Differences in analytical sensitivity were addressed by treating concentrations below the limit of detection as left-censored observations using multiple imputation, a recommended approach for environmental concentration data with high proportions of non-detects and preferable to substituting fixed fractions of the detection limit ^30–32^. Imputation had its largest influence in 1998–2000, when imputed concentrations contributed 74% of estimated toxic pressure, compared with 29% in 2013 and at most 7% in 2018–2019 and 2021. To reduce bias, the imputation model included pesticide-class and active-substance effects and the hierarchical structure of sites, campaigns and samples, while estimates were pooled across imputations to propagate imputation uncertainty. Moreover, the biological response in 1998–2000 was consistent with the exposure estimate, and the key finding that parathion dominated toxic pressure in 1998–2000 rests on frequently detected concentrations rather than imputation. Together, these steps improved comparability among campaigns and reduced the influence of analytical and sampling artefacts on the estimated temporal patterns.

As some of the detected substances that were no longer authorised as plant protection products, including fipronil, permethrin and imidacloprid, are also used in biocidal products or veterinary medicines, these detections cannot be attributed unambiguously to previous agricultural use. However, the predominantly agricultural catchments and low urban land use make such sources unlikely to dominate the overall exposure pattern.

This 23-year record is, to our knowledge, unique in Europe in combining event-driven pesticide exposure measurements with biological assessments over such an extended period. Routine monitoring under the Water Framework Directive relies largely on grab samples and covers only a fraction of relevant active substances ^15^; neonicotinoids, for example, were proposed as priority substances for surface waters only in 2022, when four of the five proposed substances were no longer approved. Given these limitations, our record provides some of the most informative field-based evidence currently available for evaluating long-term ecological outcomes and can serve as a sentinel for the EU pesticide regulatory system.

### Official indicators contrast with field evidence

The absence of ecological improvement contrasts with the progress indicated by the EU Harmonised Risk Indicator 1 (HRI1), used by the European Commission and Member States to track progress in reducing pesticide use and risk. HRI1 weights annual active-substance sales according to regulatory classification and approval status, assigning lower weighting factors to low-risk or approved substances and higher factors to candidates for substitution or non-approved substances. In Germany, HRI1 declined by a factor of 1.4 between our 2013 and 2021 campaigns and by a factor of

1.7 by 2024, the latest available reporting year; across the EU, it declined by a factor of 2.3 between 2013 and 2024 ^13^. However, because HRI1 is based on the mass of active substances sold rather than their toxicity, it can favour highly potent substances that achieve efficacy at low application rates. Moreover, the high weighting of non-approved substances means that their withdrawal inherently reduces the indicator in retrospective comparisons.

Our data cannot formally exclude a decline in toxic pressure between 2013 and 2021, but our estimates indicate, at best, only a small ecological improvement: according to the field exposure– effect relationship, the estimated change in log TU_sum_ corresponds to an increase in SPEAR_pesticides_ of 0.04 (95% CI: −0.09 to +0.17), well below the 0.2 units corresponding to one ecological status class; observed SPEAR_pesticides_ values did not indicate an improvement either (0.56 in 2013, 0.43 in 2021). Starting from the observed 2021 baseline, reaching good status (SPEAR_pesticides_ = 0.6) in the average stream would require a 9.1-fold reduction in toxic pressure.

### Closing the loop between pesticide regulation and field evidence

Our findings have three implications for EU pesticide regulation and monitoring. First and most fundamentally, the success of pesticide regulation must be measured where protection is intended: in the field. Approval and authorisation decisions rest on prospective predictions of exposure and effects, but whether these hold is not systematically verified ^33^. The consistent, substance-independent field exposure–effect relationship shown here provides a yardstick for such verification. We propose that event-driven monitoring of pesticide exposure in small agricultural streams using analytical methods with detection limits in ecologically relevant ranges, combined with invertebrate community assessment such as SPEAR_pesticides_, becomes a mandatory element of post-authorisation evaluation, with defined triggers for reassessment. The legal basis largely exists: Article 21(1) of Regulation (EC) No 1107/2009 allows the Commission to review an approval at any time and requires it to consider Member State requests for review ‘in the light of new scientific and technical knowledge and monitoring data’, explicitly in cases in which the objectives of the Water Framework Directive are compromised ^6^. What is missing is systematic field monitoring that could trigger such reviews, and indicators of regulatory success expressed in units of ecological effect rather than weighted sales.

Second, reassessment must be timely. Extensions averaging more than six years, for substances of which almost half ultimately failed renewal, show that the current reassessment process is already too slow. Replacing periodic renewal with unlimited approval, as currently proposed by the European Commission under the EU Omnibus X package, would remove the main mechanism by which substances that no longer meet current approval criteria are identified and eventually withdrawn ^12,34^.

Third, because substitution alone does not reduce toxic pressure, substantial reductions in pesticide use and broader implementation of effective mitigation measures will be required. Considerable reductions in pesticide use can be achieved in many cases without losses in productivity or profitability ^35^, while vegetated buffer strips, constructed wetlands and adjusted application timing can reduce pesticide inputs to surface waters ^36^. Without a feedback loop from the field to approval and authorisation, regulation will continue to change which chemicals impair stream ecosystems without changing how much.

## Methods

### Study design and data compilation

We compiled data from four monitoring campaigns that assessed runoff-associated pesticide exposure and pesticide effects on invertebrate communities in lowland streams in agricultural landscapes in Germany between 1998 and 2021. The dataset comprises 152 unique sites: 20 sampled in 1998–2000 ^16^, 23 in 2013 ^17^, 124 in 2018–2019 ^5^ and 14 in 2021 ^18^. Twenty-one sites were sampled in more than one campaign (15 in two, 4 in three and 2 in all four campaigns). Sites covered broad gradients of agricultural land use, pesticide exposure and ecological condition. An overview of monitoring-site locations and catchment land use across all campaigns is provided in Supplementary Figure 1 and Supplementary Table 1.

### Water sampling and chemical analysis

All campaigns assessed pesticide exposure during the main application period, generally between April and July, when the highest pesticide concentrations in agricultural streams are expected ^37^. Event-driven water sampling was applied in all campaigns to capture runoff-associated pesticide peaks following rainfall, using water-level- or conductivity-triggered automated samplers or event-triggered bottle samplers. Comparisons between event-driven sampling systems showed no significant differences in measured concentrations ^16^, although variation in sampling duration may affect temporal integration and thus the height of substance-specific peaks ^38^. Event-driven sampling was complemented by grab sampling in the 2018–2019 and 2021 campaigns. Target substances were selected on the basis of regional or national use or sales, expected toxicity to aquatic invertebrates and analytical feasibility, to capture substances of highest environmental relevance in each campaign. In total, the compiled dataset comprises 126 pesticides, including 29 insecticides, 39 fungicides and 58 herbicides, and 39 metabolites/plant growth regulators. Pesticide concentrations were determined by targeted gas or liquid chromatography with mass-spectrometric or compound-specific detection. Detailed descriptions of sampling, sample preparation and analytical procedures are provided in the corresponding original publications ^5,16–18^. Campaign-specific lists of analysed substances with detection limits are provided in Supplementary Table 2.

### Invertebrate sampling

Stream invertebrates were sampled several times during the main application period; to reflect effects of same-year exposure, we included only samples collected towards the end of this period, generally in June. Benthic macroinvertebrates were collected from multiple locations within a defined 50-m stream reach using Surber or kick sampling following standardised protocols. Collected organisms were preserved in ethanol and identified in the laboratory, generally to family or species level. Further details on sampling procedures and taxonomic identification are described in the corresponding original publications ^5,16–18^.

### Toxic units

To ensure comparability among campaigns, we recalculated toxic units (TU) for all datasets as

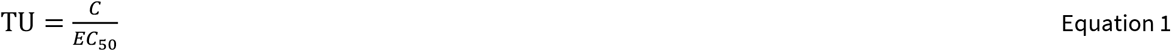

where C is the measured or imputed concentration of a substance and EC50 its acute median effect concentration for aquatic invertebrates. We compiled acute EC50 values from the US EPA ECOTOX database ^39^ for Daphnia magna and Chironomus sp. with mortality or immobility as endpoints, calculated geometric means where several values were available for the same organism, and used the EC50 of the more sensitive organism; data gaps were filled from the Pesticide Properties DataBase ^40^. The EC50 values used for toxic unit calculations are provided for each substance in Supplementary Table 3. For each sample, we calculated TU_sum_ as the sum of all substance-specific TUs, representing mixture toxicity under concentration addition. Peak toxic pressure per site and year was characterised by the maximum TU_sum_ across samples; for sites sampled in several years within a campaign, we averaged these values. To identify toxicity drivers, we determined the relative contribution of each substance to TU_sum_. On average across sites and campaigns, the largest single-substance contribution accounted for 76% (range across imputations 75–77%) of TU_sum_. We excluded 8 site-year observations (2 in 1998–2000; 2 in 2013; 4 in 2018–2019) corresponding to 2 sites (2 sites in 2013) in which event-driven samplers failed to capture runoff events (Supplementary Table 1).

### Non-detects

Concentrations below the limit of detection (LoD) made up 85.7%, 83.1%, 72.4% and 74.8% of substance–sample combinations in 1998–2000, 2013, 2018–2019 and 2021. Because highly potent substances may contribute to toxicity even below the LoD, and to avoid bias from campaign-specific analytical limits, we imputed non-detects before calculating toxic units. Multiple imputation provides an appropriate approach for left-censored concentration data with substantial proportions of non-detects, whereas substitution with fixed fractions of the LoD can bias estimates, particularly when non-detects are frequent ^30–32^. We used a Bayesian left-censored lognormal mixed-effects model fitted with the R package brms (version 2.23.0) ^41^, as it allows left-censored observations to be modelled while accounting for the hierarchical structure of the data. We treated concentrations below the LoD as left-censored observations at their observation-specific LoD. Pesticide substance and class identity were included as random intercepts to allow partial pooling, particularly for substances with few or no detections, and site and sample identity to account for the hierarchical sampling structure; exploratory analyses of detected concentrations indicated positive within-site (intraclass correlation 0.35) and within-sample correlation (0.41). The final model was concentration | cens(cens) ∼ 1 + (1 | class) + (1 | sub) + (1 | siteID) + (1 | sampleID) with a lognormal response distribution; the sample identifier combines campaign, site and sampling date. We ran four Markov chain Monte Carlo chains with 2,000 iterations each, including 1,000 warm-up iterations, resulting in 4,000 post-warm-up draws. Convergence was assessed using the potential scale reduction factor (R-hat) and effective sample sizes. All R-hat values were 1.00, with Bulk ESS ≥ 872 and Tail ESS ≥ 1,472. No divergent transitions or maximum treedepth exceedances occurred. From the fitted model we generated 100 imputed datasets, a number chosen based on the stability of estimates and their uncertainty in the subsequent analyses; for each dataset, concentrations below the LoD were sampled from the posterior predictive distribution conditional on being below the respective LoD, using the R package truncnorm (version 1.09) ^42^ on the log-concentration scale. This procedure propagates uncertainty in both the model parameters and the individual concentrations below the LoD. Substances not analysed in a campaign were not imputed. Imputed values contributed 74.1%, 29.0%, 0.9% and 6.5% of TU_sum_ in the four campaigns; contributions per insecticide class and regulatory status and their variation among imputations are given in Supplementary Tables 4 and 5.

### SPEAR_pesticides_ indicator

To assess pesticide-related ecological effects, we used the SPEAR_pesticides_ indicator, which quantifies the relative abundance of pesticide-vulnerable taxa in stream macroinvertebrate communities ^16,17^. Taxa are classified as vulnerable or invulnerable to pesticide exposure based on ecological and toxicological traits, including physiological sensitivity, generation time, exposure probability during the main pesticide application period and recolonisation potential. Because the taxon classification has been revised since the earliest campaigns, we recalculated SPEAR_pesticides_ from the compiled abundance data using a consistent classification as

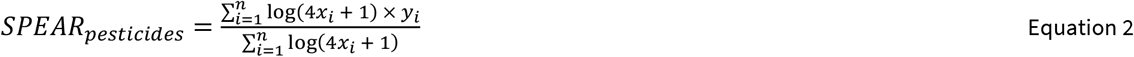

where n is the number of taxa in a sample, xi is the abundance of taxon i, and y_i_ is the vulnerability classification of taxon _i_, with y_i_ = 1 for pesticide-vulnerable taxa and y_i_ = 0 for pesticide-invulnerable taxa. The log transformation reduces the influence of taxa with mass occurrences; higher values indicate a higher proportion of vulnerable taxa and thus lower pesticide-related impairment. SPEAR_pesticides_ responds primarily to pesticide toxic pressure and is relatively independent of other environmental stressors ^5^. It has been applied across Europe ^5,16–18,43,44^, in Siberia ^45^, North and South America ^28,46,47^, Australia ^48^ and East Africa ^29^, and its relationship with toxic units did not differ among field studies from Europe, Siberia and Australia ^19^.

To harmonise SPEAR_pesticides_ across campaigns and account for differences in sampling conditions, taxonomic resolution and large-scale environmental factors unrelated to pesticide exposure (for example climate) we normalised indicator values using campaign-specific reference estimates. For each campaign, we fitted a linear exposure–effect model relating SPEAR_pesticides_ to log TU_sum_ within each imputed dataset, pooled the coefficients using Rubin’s rules ^49^ and predicted the reference value at log TU_sum_ = −5, a level of toxic pressure at which no effects on invertebrate communities are expected. Each observed SPEAR_pesticides_ value was then divided by the respective reference value. The reference values were 0.33 (1998–2000), 0.33 (2013), 0.35 (2018–2019) and 0.28 (2021). Because the reference values were similar among campaigns and lowest in 2021, normalisation is unlikely to have masked an improvement. Normalised values close to 1 indicate communities comparable to the reference condition, while lower values indicate a reduced proportion of pesticide-vulnerable taxa. Normalised values were classified with the boundaries 0.8, 0.6, 0.4 and 0.2 (high, good, moderate, poor, bad) used by Liess et al. ^5^; one status class thus spans 0.2 units. For sites sampled in several years within a campaign, we averaged SPEAR_pesticides_. 32 site-year observations (4 in 1998– 2000, 24 in 2018–2019, 4 in 2021) corresponding to 29 sites (2 in 1998–2000, 23 in 2018–2019, 4 in 2021) affected by low-flow conditions or drying were omitted from further analysis (Supplementary Table 1).

### Regulatory status

We linked the monitoring data to EU-level information on active-substance approval and national information on plant protection product authorisation, to evaluate how regulatory action was related to changes in pesticide exposure and ecological effects. We reviewed the lists of active substances authorised for use in plant protection products in the annexes to Council Directive 91/414/EEC and Regulation (EC) No 1107/2009, as well as the German database on authorised plant protection products, to determine the regulatory status of the respective active substances during each monitoring campaign. Substances were classified as authorised, authorised under extended approval (original approval period expired, approval extended under Article 17) or not authorised at the time of sampling. Substances that could still legally be used during the year of monitoring, including those subject to a grace period following the expiry or withdrawal of authorisation, were classified as authorised. In addition, we extracted approval extensions, their cumulative duration and the outcome of renewal decisions, for all measured active substances covered by Regulation (EC) No 1107/2009.

### Statistical analyses

To assess changes in toxic pressure and pesticide-related effects across campaigns, we fitted linear mixed-effects models using the lmer function from the R package lme4 (version 2.0.1) ^50^. We modelled log TU_sum_ and SPEAR_pesticides_ as response, with campaign and the percentage of cropland in the catchment as fixed effects and site identity as a random intercept (log TU_sum_ ∼ campaign + cropland + (1 | site); SPEAR_pesticides_ ∼ campaign + cropland + (1 | site)). Agricultural land accounts for differences in land-use intensity among campaigns; the random intercept accounts for repeated sampling of sites across campaigns. For log TU_sum_, models were fitted to each imputed dataset and pooled using Rubin’s rules with the R package mice (version 3.19.0) ^51^. Campaign-specific adjusted means for both response variables were estimated using estimated marginal means with the R package emmeans (version 2.0.4) ^52^. The campaign effect was tested with a pooled Wald test for log TU_sum_ and a joint F-test for SPEAR_pesticides_. Annual changes were estimated from the adjusted means with linear contrasts, assigning each campaign its midpoint year (1999, 2013, 2018.5 and 2021). The 2013–2021 change was estimated as a linear contrast between the corresponding campaign-specific adjusted means. Because community effects scale with the logarithm of toxic pressure, changes in TU_sum_ were translated into the corresponding change in SPEAR_pesticides_ using the average slope of the exposure–effect relationship of −0.175 SPEAR_pesticides_ units per log TU_sum_ unit. We regarded a change of 0.2 SPEAR_pesticides_ units, the width of one ecological status class, as ecologically relevant.

To test whether exposure–effect relationships differed among campaigns, we fitted linear mixed-effect models with SPEAR_pesticides_ as response, log TU_sum_, campaign and their interaction as fixed effects and site identity as a random intercept (SPEAR_pesticides_ ∼ log TU_sum_ × campaign + (1 | site)), pooled across imputations as above, and compared the full model with a model without the interaction using a pooled Wald test. Campaign-specific slopes and 95% confidence intervals were estimated using the emtrends function from the R package emmeans (version 2.0.4) ^52^ and pooled across imputations using Rubin’s rules. Fixed-effects regression lines and 95% confidence intervals were obtained from pooled predictions using the R package marginaleffects (version 0.32.0) ^53^. Marginal R^2^ was calculated for each imputed model using the Nakagawa R^2^ approach implemented in the R package performance (version 0.17.1) ^54^, and estimates were pooled across imputations using Rubin’s rules. Residuals were checked visually for normality and homoscedasticity using diagnostic plots produced with the R package performance (version 0.17.1) ^54^.

To assess shifts in toxicity drivers, we calculated the relative contribution of individual pesticides to TU_sum_. For each imputed dataset, contributions were derived at the site level from the sample defining the site-specific maximum TU_sum_, averaged across sites within each campaign and then across imputed datasets. Contributions were summarised by pesticide class and by regulatory status at the time of sampling. To focus on toxicity patterns in agricultural streams, this analysis was restricted to sites with ≥20% cropland in their catchments. All analyses were performed in R (version 4.6.1) ^55^; figures were produced with ggplot2 (version 4.0.3) ^56^.

### Sensitivity analyses

To assess robustness to differences in site composition among campaigns, we repeated the campaign-comparison and temporal-trend analyses described above for the subset of 21 sites sampled in more than one campaign, using the same model specifications (Supplementary Figure 2). To assess sensitivity to imputation, we quantified the contribution of imputed concentrations to TU_sum_ by campaign, insecticide class and regulatory status, and summarised variation in class- and status-specific contributions across imputations using the 2.5th and 97.5th percentiles (Supplementary Tables 4 and 5).

### Use of large language models

During the preparation of this manuscript, the authors used ChatGPT (GPT-5.5 Thinking, OpenAI) and Claude (Opus 5.5, Anthropic) to improve the structure, language, and readability. These tools were not used to formulate research hypotheses, conduct or design analyses, interpret results, draw scientific conclusions, generate or fabricate data or citations, or otherwise replace the authors’ scientific judgment. All AI-assisted content was critically reviewed by the authors and revised as necessary. The authors assume full responsibility for the accuracy, integrity, and content of the final manuscript.

## Data availability

Data will be deposited in an appropriate public repository upon publication.

## Code availability

R code will be deposited in an appropriate public repository upon publication.

## Author contributions

J.G. conceptualised the study, compiled and analysed the data, produced the figures and wrote the original draft. M.L. conceptualised the study, acquired funding and supervised the work. Both authors reviewed and edited the manuscript.

