## Supporting Information for "Two decades of EU pesticide substitution have not reduced impacts on stream invertebrates"

#### Contents

Supplementary Figures ........................................................................................................... 2

Supplementary Tables ............................................................................................................ 3

#### Supplementary Figures

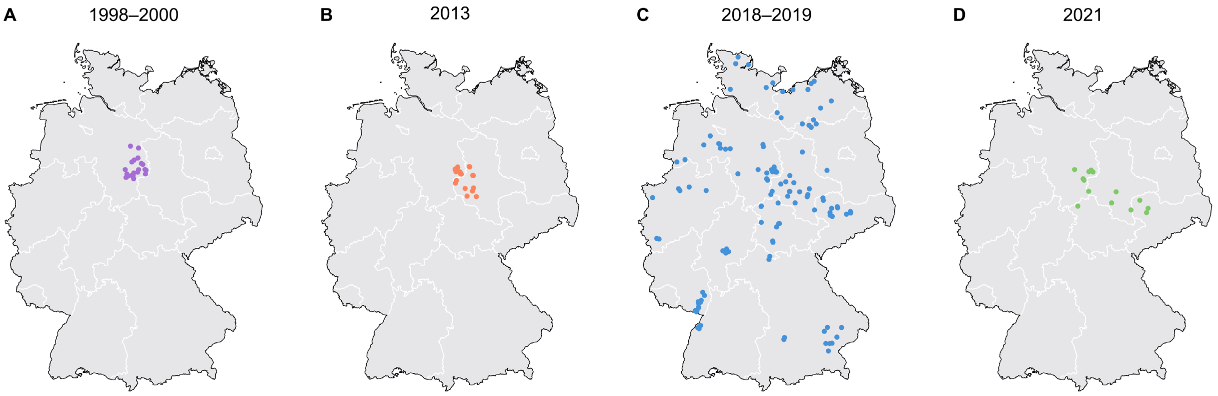


**Supplementary Figure 1: Locations of the monitoring sites.** Stream monitoring sites in Germany sampled during the four campaigns: a, 1998–2000 (n = 20); b, 2013 (n = 23); c, 2018–2019 (n = 124); d, 2021 (n = 14).

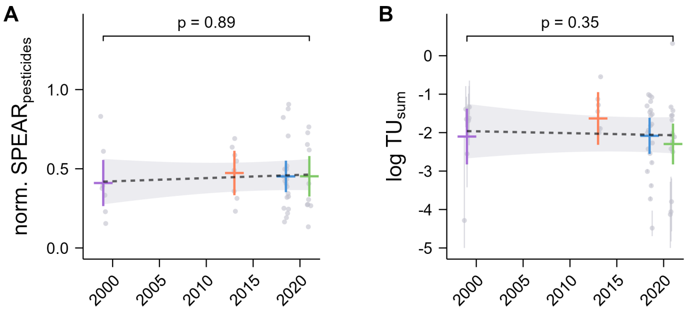


**Supplementary Figure 2: Campaign comparison for the 21 sites sampled in more than one campaign.** Campaign-specific estimated marginal means and 95% confidence intervals of normalised $SPEAR_{pesticides}$ (a) and log $TU_{sum}$ (b); dashed lines and grey bands show the linear temporal trend and its 95% confidence interval; p values refer to the campaign effect. Grey points show site-level observations; grey lines indicate the 2.5th–97.5th percentile range of log $TU_{sum}$ across imputations. n = 7, 8, 20 and 10 sites (a) and 7, 6, 21 and 14 sites (b) in 1998–2000, 2013, 2018–2019 and 2021, respectively.

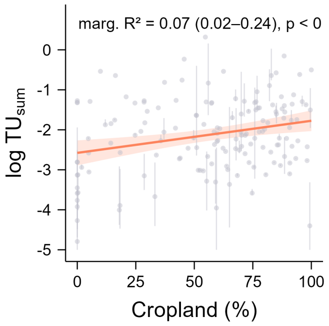


**Supplementary Figure 3: Toxic pressure increased with the proportion of cropland in the catchment.** Relationship between the proportion of cropland in the catchment and log $TU_{sum}$; line and band show fixed-effect prediction and its 95% confidence interval; grey points show site-level observations; grey lines indicate the 2.5th–97.5th percentile range of log $TU_{sum}$ across imputations.

### Supplementary Tables

**Supplementary Table 1: Overview of monitoring sites, catchment sizes and land use.** Superscript a indicates sites excluded from log $TU_{sum}$ analyses; superscript b indicates sites excluded from $SPEAR_{pesticides}$ analyses.

| Campaign | Site | Catchment ($km^2$) | Cropland (%) | Pastures (%) | Forest (%) | Urban (%) |
|---|---|---|---|---|---|---|
| 1998–2000 | S1 | 5.6 | 0 | 0 | 100 | 0 |
| 1998–2000 | S2 | 10.8 | 82 | 0 | 14 | 4 |
| 1998–2000 | S3 | 11.6 | 100 | 0 | 0 | 0 |
| 1998–2000 | S4 | 9.0 | 51 | 0 | 46 | 3 |
| 1998–2000 | S5 | 16.1 | 83 | 0 | 15 | 2 |
| 1998–2000 | S6 | 11.7 | 56 | 3 | 38 | 3 |
| 1998–2000 | S7 | 18.5 | 76 | 11 | 8 | 5 |
| 1998–2000 | S8 | 11.5 | 65 | 1 | 33 | 1 |
| 1998–2000 | S9 [b] | 15.7 | 70 | 0 | 30 | 0 |
| 1998–2000 | S10 [b] | 25.9 | 59 | 1 | 34 | 4 |
| 1998–2000 | S11 | 18.5 | 71 | 4 | 20 | 4 |
| 1998–2000 | S12 | 10.4 | 77 | 0 | 20 | 3 |
| 1998–2000 | S13 | 13.0 | 68 | 5 | 26 | 1 |
| 1998–2000 | S14 | 7.5 | 96 | 0 | 0 | 4 |
| 1998–2000 | S15 | 6.7 | 37 | 4 | 59 | 0 |
| 1998–2000 | S16 | 11.4 | 86 | 0 | 11 | 3 |
| 1998–2000 | S17 | 8.5 | 87 | 0 | 10 | 3 |
| 1998–2000 | S18 | 16.7 | 70 | 0 | 24 | 6 |
| 1998–2000 | S19 | 22.0 | 20 | 0 | 80 | 0 |
| 1998–2000 | S20 | 5.5 | 88 | 0 | 12 | 0 |
| 2013 | S21 | 14.9 | 12 | 0 | 67 | 21 |
| 2013 | S22 | 174.8 | 29 | 12 | 56 | 2 |
| 2013 | S23 | 5.7 | 59 | 0 | 41 | 0 |

| | | | | | | | |
|---|---|---|---|---|---|---|---|
| 2013 | S24 | | 1.9 | 69 | 0 | 31 | 0 |
| 2013 | S25 | | 5.4 | 36 | 5 | 17 | 41 |
| 2013 | S26 | | 290.5 | 33 | 7 | 54 | 6 |
| 2013 | S27 | | 5.8 | 95 | 0 | 0 | 5 |
| 2013 | S28 | | 3.4 | 91 | 0 | 0 | 9 |
| 2013 | S29 | | 8.3 | 62 | 2 | 24 | 12 |
| 2013 | S30 | | 7.5 | 5 | 0 | 95 | 0 |
| 2013 | S31 | [a] | 6.6 | 0 | 0 | 100 | 0 |
| 2013 | S14 | | 7.5 | 96 | 0 | 0 | 4 |
| 2013 | S16 | | 11.4 | 85 | 0 | 12 | 3 |
| 2013 | S18 | | 16.7 | 67 | 1 | 25 | 8 |
| 2013 | S32 | | 3.2 | 76 | 0 | 10 | 13 |
| 2013 | S33 | | 19.4 | 25 | 7 | 66 | 2 |
| 2013 | S34 | | 141.7 | 50 | 5 | 36 | 8 |
| 2013 | S35 | | 1.5 | 99 | 0 | 1 | 0 |
| 2013 | S36 | | 1.6 | 66 | 0 | 21 | 13 |
| 2013 | S20 | [a] | 5.5 | 89 | 0 | 11 | 0 |
| 2013 | S37 | | 8.0 | 55 | 13 | 25 | 6 |
| 2013 | S38 | | 68.2 | 18 | 16 | 63 | 2 |
| 2013 | S39 | | 12.7 | 16 | 10 | 71 | 3 |
| 2018–2019 | S40 | | 15.2 | 57 | 15 | 26 | 1 |
| 2018–2019 | S41 | | 15.0 | 87 | 6 | 7 | 0 |
| 2018–2019 | S42 | | 5.8 | 72 | 8 | 18 | 1 |
| 2018–2019 | S43 | [b] | 17.6 | 64 | 20 | 14 | 1 |
| 2018–2019 | S44 | | 10.7 | 78 | 13 | 6 | 4 |
| 2018–2019 | S45 | | 17.7 | 87 | 9 | 2 | 2 |
| 2018–2019 | S46 | | 10.2 | 29 | 18 | 39 | 15 |

| | | | | | | | |
|---|---|---|---|---|---|---|---|
| 2018–2019 | S47 | | 15.3 | 29 | 7 | 60 | 5 |
| 2018–2019 | S48 | | 6.4 | 58 | 19 | 3 | 15 |
| 2018–2019 | S49 | | 18.5 | 60 | 8 | 24 | 8 |
| 2018–2019 | S50 | | 19.5 | 61 | 3 | 33 | 4 |
| 2018–2019 | S51 | | 5.4 | 80 | 16 | 4 | 0 |
| 2018–2019 | S52 | | 15.3 | 0 | 0 | 100 | 0 |
| 2018–2019 | S53 | | 6.2 | 74 | 10 | 15 | 0 |
| 2018–2019 | S54 | | 6.1 | 75 | 11 | 0 | 14 |
| 2018–2019 | S55 | | 12.4 | 79 | 6 | 12 | 3 |
| 2018–2019 | S56 | | 14.2 | 34 | 38 | 27 | 2 |
| 2018–2019 | S57 | | 12.0 | 62 | 22 | 14 | 1 |
| 2018–2019 | S58 | | 9.6 | 72 | 28 | 0 | 0 |
| 2018–2019 | S59 | | 27.9 | 77 | 5 | 16 | 2 |
| 2018–2019 | S60 | | 11.9 | 72 | 3 | 23 | 2 |
| 2018–2019 | S61 | | 17.1 | 75 | 3 | 17 | 5 |
| 2018–2019 | S62 | | 8.1 | 14 | 38 | 45 | 3 |
| 2018–2019 | S63 | | 11.8 | 44 | 15 | 38 | 4 |
| 2018–2019 | S64 | | 17.9 | 30 | 17 | 50 | 3 |
| 2018–2019 | S65 | | 23.6 | 76 | 2 | 10 | 11 |
| 2018–2019 | S66 | | 6.2 | 57 | 0 | 42 | 0 |
| 2018–2019 | S67 | [b] | 11.8 | 70 | 17 | 13 | 0 |
| 2018–2019 | S68 | | 9.2 | 47 | 13 | 39 | 0 |
| 2018–2019 | S69 | | 22.0 | 70 | 16 | 13 | 0 |
| 2018–2019 | S70 | [b] | 10.4 | 79 | 16 | 4 | 0 |
| 2018–2019 | S71 | | 9.9 | 93 | 6 | 0 | 1 |
| 2018–2019 | S72 | | 12.8 | 73 | 7 | 19 | 0 |
| 2018–2019 | S73 | [b] | 14.9 | 64 | 22 | 14 | 0 |

| | | | | | | | |
|---|---|---|---|---|---|---|---|
| 2018–2019 | S74 | | 14.1 | 44 | 23 | 31 | 2 |
| 2018–2019 | S75 | | 15.1 | 55 | 10 | 34 | 2 |
| 2018–2019 | S30 | [b] | 7.5 | 5 | 0 | 95 | 0 |
| 2018–2019 | S31 | | 6.6 | 0 | 0 | 100 | 0 |
| 2018–2019 | S14 | | 7.5 | 96 | 0 | 0 | 4 |
| 2018–2019 | S76 | [b] | 9.8 | 85 | 8 | 6 | 0 |
| 2018–2019 | S15 | | 6.7 | 37 | 0 | 63 | 0 |
| 2018–2019 | S77 | | 15.9 | 91 | 4 | 6 | 0 |
| 2018–2019 | S16 | | 11.4 | 85 | 0 | 12 | 3 |
| 2018–2019 | S78 | | 8.8 | 54 | 5 | 34 | 7 |
| 2018–2019 | S79 | | 16.4 | 63 | 11 | 22 | 4 |
| 2018–2019 | S80 | | 4.5 | 73 | 0 | 27 | 0 |
| 2018–2019 | S17 | | 8.5 | 84 | 3 | 10 | 3 |
| 2018–2019 | S18 | | 16.7 | 67 | 1 | 25 | 8 |
| 2018–2019 | S81 | | 31.5 | 0 | 0 | 100 | 0 |
| 2018–2019 | S82 | [b] | 12.7 | 92 | 0 | 4 | 4 |
| 2018–2019 | S83 | | 16.4 | 88 | 6 | 7 | 0 |
| 2018–2019 | S84 | | 26.0 | 79 | 3 | 14 | 3 |
| 2018–2019 | S85 | [b] | 4.8 | 91 | 9 | 0 | 0 |
| 2018–2019 | S32 | | 3.2 | 76 | 0 | 10 | 13 |
| 2018–2019 | S19 | | 22.0 | 16 | 0 | 84 | 0 |
| 2018–2019 | S86 | | 22.2 | 68 | 7 | 25 | 0 |
| 2018–2019 | S87 | | 2.4 | 0 | 0 | 100 | 0 |
| 2018–2019 | S88 | | 19.8 | 81 | 8 | 9 | 3 |
| 2018–2019 | S33 | | 19.4 | 25 | 7 | 66 | 2 |
| 2018–2019 | S89 | | 1.8 | 0 | 0 | 100 | 0 |
| 2018–2019 | S90 | | 16.0 | 78 | 19 | 0 | 3 |

| | | | | | | | |
|---|---|---|---|---|---|---|---|
| 2018–2019 | S91 | | 16.8 | 81 | 11 | 8 | 0 |
| 2018–2019 | S92 | b | 17.9 | 60 | 11 | 23 | 5 |
| 2018–2019 | S93 | | 5.4 | 84 | 10 | 0 | 6 |
| 2018–2019 | S94 | b | 4.9 | 100 | 0 | 0 | 0 |
| 2018–2019 | S95 | | 5.0 | 89 | 9 | 1 | 1 |
| 2018–2019 | S96 | | 20.5 | 57 | 8 | 31 | 4 |
| 2018–2019 | S97 | | 2.1 | 25 | 0 | 75 | 0 |
| 2018–2019 | S98 | | 5.8 | 14 | 0 | 81 | 4 |
| 2018–2019 | S99 | | 3.7 | 0 | 0 | 100 | 0 |
| 2018–2019 | S100 | b | 70.4 | 5 | 0 | 89 | 6 |
| 2018–2019 | S101 | | 29.2 | 42 | 6 | 49 | 3 |
| 2018–2019 | S102 | | 29.6 | 16 | 11 | 65 | 7 |
| 2018–2019 | S103 | | 10.4 | 21 | 0 | 63 | 16 |
| 2018–2019 | S104 | | 18.4 | 46 | 6 | 32 | 17 |
| 2018–2019 | S105 | | 19.9 | 29 | 2 | 66 | 3 |
| 2018–2019 | S106 | | 10.7 | 0 | 3 | 97 | 0 |
| 2018–2019 | S107 | | 2.1 | 0 | 0 | 100 | 0 |
| 2018–2019 | S108 | | 7.5 | 35 | 0 | 59 | 6 |
| 2018–2019 | S109 | b | 3.2 | 0 | 0 | 100 | 0 |
| 2018–2019 | S110 | | 5.8 | 37 | 0 | 57 | 6 |
| 2018–2019 | S111 | | 2.2 | 0 | 0 | 100 | 0 |
| 2018–2019 | S112 | | 10.7 | 10 | 0 | 90 | 0 |
| 2018–2019 | S113 | b | 8.9 | 54 | 44 | 1 | 1 |
| 2018–2019 | S114 | | 8.6 | 56 | 29 | 16 | 0 |
| 2018–2019 | S115 | | 18.8 | 83 | 11 | 2 | 5 |
| 2018–2019 | S116 | | 13.0 | 88 | 9 | 3 | 0 |
| 2018–2019 | S117 | | 16.5 | 51 | 15 | 31 | 4 |

| | | | | | | |
|---|---|---|---|---|---|---|
| 2018–2019 | S118 | 5.1 | 52 | 44 | 4 | 0 |
| 2018–2019 | S119 [b] | 7.0 | 5 | 0 | 95 | 0 |
| 2018–2019 | S120 | 9.5 | 98 | 0 | 1 | 0 |
| 2018–2019 | S121 [b] | 5.4 | 5 | 0 | 95 | 0 |
| 2018–2019 | S122 | 18.9 | 83 | 10 | 7 | 0 |
| 2018–2019 | S123 | 12.9 | 85 | 0 | 0 | 15 |
| 2018–2019 | S124 | 12.4 | 60 | 10 | 23 | 4 |
| 2018–2019 | S125 | 17.1 | 70 | 16 | 10 | 4 |
| 2018–2019 | S126 | 29.0 | 51 | 8 | 36 | 5 |
| 2018–2019 | S127 [b] | 3.7 | 18 | 0 | 82 | 0 |
| 2018–2019 | S128 | 14.5 | 94 | 5 | 0 | 1 |
| 2018–2019 | S129 [b] | 13.7 | 98 | 1 | 0 | 1 |
| 2018–2019 | S130 | 4.1 | 45 | 18 | 36 | 1 |
| 2018–2019 | S131 | 31.4 | 94 | 3 | 0 | 3 |
| 2018–2019 | S132 | 14.3 | 89 | 0 | 0 | 11 |
| 2018–2019 | S133 [b] | 263.9 | 55 | 3 | 32 | 10 |
| 2018–2019 | S134 | 13.1 | 86 | 0 | 10 | 4 |
| 2018–2019 | S135 | 14.8 | 88 | 1 | 4 | 6 |
| 2018–2019 | S136 [b] | 34.5 | 92 | 3 | 2 | 3 |
| 2018–2019 | S137 | 1.2 | 0 | 0 | 100 | 0 |
| 2018–2019 | S138 [b] | 32.3 | 68 | 9 | 18 | 5 |
| 2018–2019 | S20 | 5.5 | 89 | 0 | 11 | 0 |
| 2018–2019 | S139 | 8.8 | 27 | 7 | 58 | 8 |
| 2018–2019 | S140 | 11.0 | 42 | 6 | 46 | 6 |
| 2018–2019 | S141 | 18.0 | 98 | 0 | 0 | 2 |
| 2018–2019 | S142 | 22.9 | 86 | 0 | 9 | 5 |
| 2018–2019 | S143 [b] | 40.3 | 91 | 1 | 0 | 7 |

| | | | | | | |
|---|---|---|---|---|---|---|
| 2018–2019 | S144 | 1.7 | 0 | 9 | 91 | 0 |
| 2018–2019 | S145 | 7.9 | 0 | 0 | 100 | 0 |
| 2018–2019 | S146 | 5.1 | 76 | 13 | 11 | 0 |
| 2018–2019 | S147 | 1.6 | 2 | 0 | 98 | 0 |
| 2018–2019 | S148 [b] | 13.2 | 53 | 11 | 36 | 0 |
| 2018–2019 | S149 | 20.3 | 60 | 18 | 20 | 3 |
| 2018–2019 | S150 [b] | 16.8 | 59 | 14 | 10 | 17 |
| 2018–2019 | S151 | 7.7 | 88 | 0 | 0 | 11 |
| 2018–2019 | S152 | 3.8 | 92 | 0 | 0 | 8 |
| 2021 | S31 [b] | 6.6 | 0 | 0 | 100 | 0 |
| 2021 | S14 [b] | 7.5 | 96 | 0 | 0 | 4 |
| 2021 | S16 | 11.4 | 85 | 0 | 12 | 3 |
| 2021 | S80 [b] | 4.5 | 73 | 0 | 27 | 0 |
| 2021 | S17 | 8.5 | 84 | 3 | 10 | 3 |
| 2021 | S89 | 1.8 | 0 | 0 | 100 | 0 |
| 2021 | S122 | 18.9 | 83 | 10 | 7 | 0 |
| 2021 | S123 [b] | 12.9 | 85 | 0 | 0 | 15 |
| 2021 | S126 | 29.0 | 51 | 8 | 36 | 5 |
| 2021 | S131 | 31.4 | 94 | 3 | 0 | 3 |
| 2021 | S134 | 13.1 | 86 | 0 | 10 | 4 |
| 2021 | S137 | 1.2 | 0 | 0 | 100 | 0 |
| 2021 | S141 | 18.0 | 98 | 0 | 0 | 2 |
| 2021 | S147 | 1.6 | 2 | 0 | 98 | 0 |

**Supplementary Table 2: Analytes included in the monitoring campaigns and corresponding limits of detection (LoD).** Blank cells indicate that a substance was not analysed in the respective campaign.

| Substance | 1998–2000 LoD [µg/L] | 2013 LoD [µg/L] | 2018–2019 LoD [µg/L] | 2021 LoD [µg/L] |
|---|---|---|---|---|
| Azoxystrobin | 0.02 | 0.001 | 0.0001 | 0.01 |
| Bixafen | | | 0.00003 | 0.001 |
| Boscalid | | 0.02 | 0.0001 | 0.005 |
| Carbendazim | | 0.001 | | |
| Cyazofamid | | 0.002 | 0.0005 | 0.005 |
| Cyflufenamid | | 0.002 | | |
| Cyprodinil | | 0.004 | 0.0005 | 0.005 |
| Difenoconazole | | 0.005 | 0.0001 | 0.001 |
| Dimethomorph | | | 0.0003 | 0.001 |
| Dimoxystrobin | | | 0.00005 | 0.0005 |
| Epoxiconazole | 0.02 | 0.005 | 0.0001 | 0.0005 |
| Fenpropidin | | 0.002 | 0.0001 | 0.0002 |
| Fenpropimorph | 0.05 | 0.005 | | |
| Fluazinam | | | 0.00005 | 0.0005 |
| Fludioxonil | | 0.002 | 0.00005 | 0.001 |
| Flusilazole | | 0.003 | | |
| (R)-flutriafol | | 0.002 | | |
| Fluxapyroxad | | | 0.00005 | 0.001 |
| Fuberidazole | | 0.001 | | |
| Imazalil | | 0.025 | | |
| Isopyrazam | | | 0.0005 | 0.001 |
| Kresoxim-methyl | 0.02 | | 0.00125 | 0.025 |
| Mandipropamid | | 0.001 | | |
| Metalaxyl | | 0.001 | | |
| Metrafenone | | 0.003 | 0.00005 | 0.0005 |

| | | | | |
|---|---|---|---|---|
| Penconazole | | 0.003 | | |
| Picoxystrobin | | 0.001 | 0.00005 | 0.001 |
| Prochloraz | | 0.005 | 0.0002 | 0.005 |
| Propamocarb | | 0.001 | 0.00005 | 0.0005 |
| Propiconazole | 0.02 | 0.005 | 0.00075 | 0.005 |
| Pyraclostrobin | | 0.002 | 0.00005 | 0.0005 |
| Silthiofam | | 0.001 | | |
| Spiroxamine | | 0.001 | 0.0001 | 0.001 |
| Tebuconazole | 0.05 | | 0.00005 | 0.001 |
| Thiophanate-methyl | | 0.001 | | |
| Tolylfluanid | | 0.02 | | |
| Triadimenol | | | 0.00175 | 0.025 |
| Trifloxystrobin | | 0.001 | 0.00005 | 0.005 |
| Triticonazole | | 0.005 | | |
| 2.4-D | | | 0.0005 | 0.1 |
| Aclonifen | | | 0.0001 | 0.005 |
| Alachlor | | 0.005 | | |
| Amidosulfuron | | 0.001 | | |
| Atrazine | | 0.001 | | |
| Bentazone | | | 0.0001 | 0.0005 |
| Bromoxynil | | | 0.0001 | 0.0005 |
| Carfentrazone-ethyl | | 0.001 | | |
| Chloridazon | 0.05 | 0.002 | 0.0002 | 0.01 |
| Chlorotoluron | | | 0.000075 | 0.025 |
| Clomazone | | 0.001 | | |
| Cybutryne | | 0.002 | | |
| Cycloxydim | | 0.01 | | |

| | | | | |
|---|---|---|---|---|
| Dichlorprop-P | | | 0.0005 | 0.01 |
| Diflufenican | | 0.01 | 0.0001 | 0.001 |
| Dimethachlor | | | 0.00007 | 0.001 |
| Dimethenamid-P | | | 0.00005 | 0.0005 |
| Diuron | | 0.002 | | |
| Ethofumesate | 0.05 | 0.003 | 0.0002 | 0.005 |
| Florasulam | | | 0.0001 | 0.001 |
| Flufenacet | | | 0.00005 | 0.001 |
| Flupyrsulfuron-methyl-sodium | | 0.001 | | |
| Fluroxypyr | | | 0.0007 | 0.025 |
| Flurtamone | | 0.001 | 0.00006 | 0.0005 |
| Foramsulfuron | | 0.005 | 0.0001 | 0.005 |
| Hexazinone | | 0.001 | | |
| Iodosulfuron-methyl | | | 0.0003 | 0.005 |
| Isoproturon | 0.05 | 0.001 | 0.000275 | 0.005 |
| Lenacil | | | 0.00015 | 0.005 |
| MCPA | | | 0.00045 | 0.025 |
| Mecoprop-P | | | 0.0002 | 0.002 |
| Mesotrione | | | 0.001 | 0.005 |
| Metamitron | 0.05 | 0.003 | 0.0005 | 0.005 |
| Metazachlor | | 0.001 | 0.00005 | 0.0005 |
| Methabenzthiazuron | | 0.001 | | |
| Metolachlor | | 0.001 | | |
| Metribuzin | 0.05 | 0.005 | 0.0002 | 0.005 |
| Napropamide | | | 0.00005 | 0.001 |
| Nicosulfuron | | | 0.00005 | 0.0005 |
| Pendimethalin | 0.02 | 0.01 | | |

| | | | | |
|---|---|---|---|---|
| Pethoxamid | | | 0.00005 | 0.0005 |
| Phenmedipham | | 0.001 | | |
| Propyzamide | | 0.001 | 0.0001 | 0.001 |
| Prosulfocarb | 0.05 | | 0.00005 | 0.0005 |
| Prosulfuron | | | 0.00005 | 0.0005 |
| Pyroxsulam | | | 0.00005 | 0.0005 |
| Quinmerac | | 0.001 | 0.00005 | 0.001 |
| Rimsulfuron | | 0.002 | | |
| S-Metolachlor | | | 0.000075 | 0.0005 |
| Simazine | | 0.001 | | |
| Sulcotrione | | | 0.0005 | |
| Sulfosulfuron | | 0.001 | | |
| Terbuthylazine | | 0.001 | 0.00005 | 0.001 |
| Terbutryn | | 0.005 | | |
| Topramezone | | | 0.0007 | 0.005 |
| Tribenuron-methyl | | 0.001 | | |
| Triflusulfuron-methyl | | 0.002 | | |
| Tritosulfuron | | | 0.00025 | 0.005 |
| Acetamiprid | | 0.001 | 0.0001 | 0.005 |
| Alpha-cypermethrin | | 0.0005 | | |
| Beta-cyfluthrin | 0.05 | 0.001 | | |
| Bifenthrin | | | | 0.00003 |
| Carbofuran | | 0.001 | | |
| Chlorantraniliprole | | | 0.0002 | 0.005 |
| Chlorfenvinphos | | 0.002 | | |
| Clothianidin | | 0.01 | 0.00015 | 0.005 |
| Cyantraniliprole | | | | 0.005 |

| | | | | |
|---|---|---|---|---|
| Cyfluthrin | | 0.001 | | |
| Cypermethrin | 0.05 | | | 0.015 |
| Deltamethrin | | 0.001 | | 0.0075 |
| Dichlorvos | | 0.01 | | |
| Dimethoate | | 0.001 | 0.00005 | 0.001 |
| Esfenvalerate | 0.02 | 0.0003 | | 0.000075 |
| Etofenprox | | | | 0.00015 |
| Fipronil | | | 2.12121E-05 | 0.0001 |
| Imidacloprid | | 0.005 | 0.0001 | 0.001 |
| Lambda-cyhalothrin | 0.05 | 0.0001 | | 0.000075 |
| Lindane | 0.02 | | | |
| Methiocarb | | 0.002 | 0.0001 | 0.001 |
| Parathion | 0.02 | | | |
| Permethrin | | | | 0.000075 |
| Pirimicarb | 0.05 | 0.001 | 0.00005 | 0.001 |
| Spinosad | | | 0.00075 | |
| Tau-fluvalinate | | 0.001 | | |
| Tefluthrin | | | | 0.000015 |
| Thiacloprid | | 0.001 | 0.00005 | 0.0007 |
| Thiamethoxam | | 0.002 | 0.00005 | 0.0005 |
| 2.6-dichlorobenzamide | | 0.002 | | |
| Aminobifenox acid | | | 0.0025 | |
| Desethylatrazine | | 0.001 | | |
| 6-deisopropyl atrazine | | 0.005 | | |
| Azoxystrobin acid | | | 0.0001 | 0.001 |
| Bifenox acid | 0.02 | | 0.0012 | |
| Tetrahydrophthalamic acid | | | 0.0005 | 0.025 |

| | | | |
|---|---|---|---|
| Tetrahydrophthalimide | | 0.00075 | 0.1 |
| Chloridazon-desphenyl | | 0.0002 | 0.025 |
| Chloridazon-methyl-desphenyl | | 0.0001 | 0.001 |
| Chlorothalonil R471811 | | 0.0005 | 0.003 |
| Desamino-metamitron | | 0.0005 | 0.005 |
| Desmethyl chlorotoluron | | 0.0003 | |
| Dimethachlor CGA 354742 | | 0.001 | 0.002 |
| Dimethachlor CGA 369873 | | 0.0001 | 0.005 |
| Dimethenamid-P M23 | | 0.0025 | 0.1 |
| Dimethenamid-P M27 | | 0.0005 | 0.005 |
| Flufenacet ESA | | 0.0005 | 0.005 |
| Flufenacet thiadone (AE 1258593) | | 0.002 | 0.025 |
| Flumioxazin APF | | 0.002 | |
| Hexythiazox | 0.005 | | |
| Metazachlor BH 479-11 | | 0.00005 | |
| Metazachlor BH 479-12 | | 0.0005 | |
| Metazachlor BH 479-9 | | 0.00025 | |
| Metazachlor ESA | | 0.0005 | 0.005 |
| Metolachlor CGA 357704 | | 0.0008 | 0.01 |
| Metolachlor CGA 368208 | | 0.0005 | 0.01 |
| Metolachlor ESA | | 0.0005 | 0.005 |
| Metolachlor NOA 413173 | | 0.0025 | 0.025 |
| Metolachlor OA | | 0.0013 | 0.1 |
| Prothioconazole-desthio (AE 1194888) | 0.003 | 0.0001 | 0.001 |
| Quinmerac BH 518-2 | | 0.003 | |
| Quizalofop free acid | | 0.00035 | 0.005 |
| Terbuthylazine-desethyl-2-hydroxy | 0.001 | 0.00075 | 0.001 |

| | | | | |
|---|---|---|---|---|
| Thiacloprid-amide | | | 0.0001 | 0.001 |
| Trifloxystrobin CGA 321113 | | | 0.0001 | 0.001 |
| Tritosulfuron BH 635-2 | | | 0.0005 | |
| Flurprimidol | | 0.002 | | |
| Trinexapac-ethyl | | 0.002 | | |

**Supplementary Table 3: EC50 values used for toxic unit calculations.** Substance type is classified as fungicide (F), herbicide (H), insecticide (I), metabolite (M) or plant growth regulator (P). CAS numbers and data sources are provided for each substance.

| Substance | Type | CAS | EC50 (µg/L) | Source |
|---|---|---|---|---|
| Azoxystrobin | F | 131860-33-8 | 3289.679 | EPA ECOTOX |
| Bixafen | F | 581809-46-3 | 1200.000 | PPDB |
| Boscalid | F | 188425-85-6 | 5330.000 | PPDB |
| Carbendazim | F | 10605-21-7 | 83.829 | EPA ECOTOX |
| Cyazofamid | F | 120116-88-3 | 413.000 | EPA ECOTOX |
| Cyflufenamid | F | 180409-60-3 | 1730.000 | PPDB |
| Cyprodinil | F | 121552-61-2 | 32.000 | EPA ECOTOX |
| Difenoconazole | F | 119446-68-3 | 53.033 | EPA ECOTOX |
| Dimethomorph | F | 110488-70-5 | 20100.000 | PPDB |
| Dimoxystrobin | F | 149961-52-4 | 39.400 | PPDB |
| Epoxiconazole | F | 135319-73-2 | 5310.333 | EPA ECOTOX |
| Fenpropidin | F | 67306-00-7 | 1803.139 | EPA ECOTOX |
| Fenpropimorph | F | 67564-91-4 | 2792.788 | EPA ECOTOX |
| Fluazinam | F | 79622-59-6 | 199.900 | EPA ECOTOX |
| Fludioxonil | F | 131341-86-1 | 900.000 | EPA ECOTOX |
| Flusilazole | F | 85509-19-9 | 1419.289 | EPA ECOTOX |
| (R)-flutriafol | F | 76674-21-0 | 67100.000 | EPA ECOTOX |
| Fluxapyroxad | F | 907204-31-3 | 6780.000 | PPDB |

| | | | | |
|---|---|---|---|---|
| Fuberidazole | F | 3878-19-1 | 4700.000 | PPDB |
| Imazalil | F | 35554-44-0 | 2674.661 | EPA ECOTOX |
| Isopyrazam | F | 881685-58-1 | 44.000 | PPDB |
| Kresoxim-methyl | F | 143390-89-0 | 383.635 | EPA ECOTOX |
| Mandipropamid | F | 374726-62-2 | 7100.000 | EPA ECOTOX |
| Metalaxyl | F | 57837-19-1 | 109612.418 | EPA ECOTOX |
| Metrafenone | F | 220899-03-6 | 920.000 | PPDB |
| Penconazole | F | 66246-88-6 | 3694.447 | EPA ECOTOX |
| Picoxystrobin | F | 117428-22-5 | 24.000 | PPDB |
| Prochloraz | F | 67747-09-5 | 3888.809 | EPA ECOTOX |
| Propamocarb | F | 24579-73-5 | 106000.000 | PPDB |
| Propiconazole | F | 60207-90-1 | 8856.995 | EPA ECOTOX |
| Pyraclostrobin | F | 175013-18-0 | 132.469 | EPA ECOTOX |
| Silthiofam | F | 175217-20-6 | 14000.000 | PPDB |
| Spiroxamine | F | 118134-30-8 | 4164.133 | EPA ECOTOX |
| Tebuconazole | F | 107534-96-3 | 1532.665 | EPA ECOTOX |
| Thiophanate-methyl | F | 23564-05-8 | 9295.160 | EPA ECOTOX |
| Tolylfluanid | F | 731-27-1 | 190.000 | EPA ECOTOX |
| Triadimenol | F | 55219-65-3 | 2500.000 | EPA ECOTOX |
| Trifloxystrobin | F | 141517-21-7 | 105.903 | EPA ECOTOX |
| Triticonazole | F | 131983-72-7 | 7600.000 | EPA ECOTOX |
| 2.4-D | H | 94-75-7 | 6538.930 | EPA ECOTOX |
| Aclonifen | H | 74070-46-5 | 1200.000 | PPDB |
| Alachlor | H | 15972-60-8 | 7027.404 | EPA ECOTOX |
| Amidosulfuron | H | 120923-37-7 | 80000.000 | PPDB |
| Atrazine | H | 1912-24-9 | 1000.000 | EPA ECOTOX |
| Bentazone | H | 25057-89-0 | 62300.000 | EPA ECOTOX |

| | | | | |
|---|---|---|---|---|
| Bromoxynil | H | 1689-84-5 | 1900.000 | PPDB |
| Carfentrazone-ethyl | H | 128639-02-1 | 9800.000 | PPDB |
| Chloridazon | H | 1698-60-8 | 132000.000 | PPDB |
| Chlorotoluron | H | 15545-48-9 | 67000.000 | PPDB |
| Clomazone | H | 81777-89-1 | 5200.000 | EPA ECOTOX |
| Cybutryne | H | 28159-98-0 | 14680.960 | EPA ECOTOX |
| Cycloxydim | H | 101205-02-1 | 70800.000 | PPDB |
| Dichlorprop-P | H | 15165-67-0 | 100000.000 | PPDB |
| Diflufenican | H | 83164-33-4 | 240.000 | PPDB |
| Dimethachlor | H | 50563-36-5 | 24000.000 | PPDB |
| Dimethenamid-P | H | 163515-14-8 | 12000.000 | EPA ECOTOX |
| Diuron | H | 330-54-1 | 4758.973 | EPA ECOTOX |
| Ethofumesate | H | 26225-79-6 | 137171.426 | EPA ECOTOX |
| Florasulam | H | 145701-23-1 | 292000.000 | PPDB |
| Flufenacet | H | 142459-58-3 | 30900.000 | PPDB |
| Flupyrsulfuron-methyl-sodium | H | 144740-54-5 | 721000.000 | PPDB |
| Fluroxypyr | H | 69377-81-7 | 100000.000 | PPDB |
| Flurtamone | H | 96525-23-4 | 13000.000 | PPDB |
| Foramsulfuron | H | 173159-57-4 | 100000.000 | PPDB |
| Hexazinone | H | 51235-04-2 | 151799.868 | EPA ECOTOX |
| Iodosulfuron-methyl | H | 144550-36-7 | 100000.000 | PPDB |
| Isoproturon | H | 34123-59-6 | 580.000 | PPDB |
| Lenacil | H | 2164-08-1 | 8400.000 | PPDB |
| MCPA | H | 94-74-6 | 190000.000 | PPDB |
| Mecoprop-P | H | 16484-77-8 | 91000.000 | PPDB |
| Mesotrione | H | 104206-82-8 | 840000.000 | EPA ECOTOX |
| Metamitron | H | 41394-05-2 | 5700.000 | PPDB |

| | | | | |
|---|---|---|---|---|
| Metazachlor | H | 67129-08-2 | 33000.000 | PPDB |
| Methabenzthiazuron | H | 18691-97-9 | 30600.000 | PPDB |
| Metolachlor | H | 51218-45-2 | 6055.694 | EPA ECOTOX |
| Metribuzin | H | 21087-64-9 | 4180.000 | EPA ECOTOX |
| Napropamide | H | 15299-99-7 | 18793.882 | EPA ECOTOX |
| Nicosulfuron | H | 111991-09-4 | 90000.000 | PPDB |
| Pendimethalin | H | 40487-42-1 | 802.496 | EPA ECOTOX |
| Pethoxamid | H | 106700-29-2 | 23000.000 | PPDB |
| Phenmedipham | H | 13684-63-4 | 14000.000 | EPA ECOTOX |
| Propyzamide | H | 23950-58-5 | 5600.000 | PPDB |
| Prosulfocarb | H | 52888-80-9 | 510.000 | PPDB |
| Prosulfuron | H | 94125-34-5 | 120000.000 | PPDB |
| Pyroxsulam | H | 422556-08-9 | 100000.000 | PPDB |
| Quinmerac | H | 90717-03-6 | 100000.000 | PPDB |
| Rimsulfuron | H | 122931-48-0 | 1000000.000 | EPA ECOTOX |
| S-Metolachlor | H | 87392-12-9 | 26000.000 | EPA ECOTOX |
| Simazine | H | 122-34-9 | 1100.000 | EPA ECOTOX |
| Sulcotrione | H | 99105-77-8 | 848000.000 | PPDB |
| Sulfosulfuron | H | 141776-32-1 | 96000.000 | PPDB |
| Terbuthylazine | H | 5915-41-3 | 21200.000 | PPDB |
| Terbutryn | H | 886-50-0 | 6874.210 | EPA ECOTOX |
| Topramezone | H | 210631-68-8 | 100000.000 | PPDB |
| Tribenuron-methyl | H | 101200-48-0 | 894000.000 | PPDB |
| Triflusulfuron-methyl | H | 126535-15-7 | 960000.000 | PPDB |
| Tritosulfuron | H | 142469-14-5 | 100000.000 | PPDB |
| Acetamiprid | I | 135410-20-7 | 5.406 | EPA ECOTOX |
| Alpha-cypermethrin | I | 67375-30-8 | 0.285 | EPA ECOTOX |

| | | | | |
|---|---|---|---|---|
| Beta-cyfluthrin | I | 1820573-27-0 | 0.094 | EPA ECOTOX |
| Bifenthrin | I | 82657-04-3 | 0.427 | EPA ECOTOX |
| Carbofuran | I | 1563-66-2 | 51.581 | EPA ECOTOX |
| Chlorantraniliprole | I | 500008-45-7 | 11.785 | EPA ECOTOX |
| Chlorfenvinphos | I | 470-90-6 | 56.847 | EPA ECOTOX |
| Clothianidin | I | 210880-92-5 | 5.297 | EPA ECOTOX |
| Cyantraniliprole | I | 736994-63-1 | 4.700 | EPA ECOTOX |
| Cyfluthrin | I | 68359-37-5 | 0.094 | EPA ECOTOX |
| Cypermethrin | I | 52315-07-8 | 9.049 | EPA ECOTOX |
| Deltamethrin | I | 52918-63-5 | 2.192 | EPA ECOTOX |
| Dichlorvos | I | 62-73-7 | 0.746 | EPA ECOTOX |
| Dimethoate | I | 60-51-5 | 566.640 | EPA ECOTOX |
| Esfenvalerate | I | 66230-04-4 | 0.280 | EPA ECOTOX |
| Etofenprox | I | 80844-07-1 | 0.570 | EPA ECOTOX |
| Fipronil | I | 120068-37-3 | 0.181 | EPA ECOTOX |
| Imidacloprid | I | 138261-41-3 | 6.548 | EPA ECOTOX |
| Lambda-cyhalothrin | I | 91465-08-6 | 0.022 | EPA ECOTOX |
| Lindane | I | 58-89-9 | 81.281 | EPA ECOTOX |
| Methiocarb | I | 2032-65-7 | 19.000 | EPA ECOTOX |
| Parathion | I | 56-38-2 | 1.741 | EPA ECOTOX |
| Permethrin | I | 52645-53-1 | 0.333 | EPA ECOTOX |
| Pirimicarb | I | 23103-98-2 | 16.908 | EPA ECOTOX |
| Spinosad | I | 168316-95-8 | 1000.000 | PPDB |
| Tau-fluvalinate | I | 102851-06-9 | 0.940 | EPA ECOTOX |
| Tefluthrin | I | 79538-32-2 | 0.070 | PPDB |
| Thiacloprid | I | 111988-49-9 | 1.131 | EPA ECOTOX |
| Thiamethoxam | I | 153719-23-4 | 47.980 | EPA ECOTOX |

| | | | | |
|---|---|---|---|---|
| 2.6-dichlorobenzamide | M | 2008-58-4 | 856000.000 | EPA ECOTOX |
| Aminobifenox acid | M | | | |
| Desethylatrazine | M | 6190-65-4 | 35600.000 | EPA ECOTOX |
| 6-deisopropyl atrazine | M | 1007-28-9 | 34500.000 | EPA ECOTOX |
| Azoxystrobin acid | M | 1185255-09-7 | | |
| Bifenox acid | M | 53774-07-5 | | |
| Tetrahydrophthalamic acid | M | 2028-12-8 | 220000.000 | PPDB |
| Tetrahydrophthalimide | M | 85-40-5 | 120000.000 | PPDB |
| Chloridazon-desphenyl | M | 6339-19-1 | 100000.000 | PPDB |
| Chloridazon-methyl-desphenyl | M | 17254-80-7 | 100000.000 | PPDB |
| Chlorothalonil R471811 | M | | | |
| Desamino-metamitron | M | | 745000.000 | PPDB |
| Desmethyl chlorotoluron | M | | 100000.000 | PPDB |
| Dimethachlor CGA 354742 | M | | 100000.000 | PPDB |
| Dimethachlor CGA 369873 | M | | | |
| Dimethenamid-P M23 | M | 380412-59-9 | 95000.000 | PPDB |
| Dimethenamid-P M27 | M | 205939-58-8 | 100000.000 | PPDB |
| Flufenacet ESA | M | 201668-32-8 | 87300.000 | PPDB |
| Flufenacet thiadone (AE 1258593) | M | 84352-75-0 | | |
| Flumioxazin APF | M | | | |
| Hexythiazox | M | 78587-05-0 | 360.000 | PPDB |
| Metazachlor BH 479-11 | M | | | |
| Metazachlor BH 479-12 | M | | | |
| Metazachlor BH 479-9 | M | | | |
| Metazachlor ESA | M | 172960-62-2 | 93800.000 | PPDB |
| Metolachlor CGA 357704 | M | | | |
| Metolachlor CGA 368208 | M | | | |

| | | | | |
|---|---|---|---|---|
| Metolachlor ESA | M | 171118-09-5 | 100000.000 | PPDB |
| Metolachlor NOA 413173 | M | | | |
| Metolachlor OA | M | 152019-73-3 | 16600.000 | PPDB |
| Prothioconazole-desthio (AE 1194888) | M | 120983-64-4 | 10000.000 | PPDB |
| Quinmerac BH 518-2 | M | 90717-07-0 | 100000.000 | PPDB |
| Quizalofop free acid | M | 76578-12-6 | 57700.000 | PPDB |
| Terbuthylazine-desethyl-2-hydroxy | M | | 15000.000 | PPDB |
| Thiacloprid-amide | M | | 103000.000 | PPDB |
| Trifloxystrobin CGA 321113 | M | 252913-85-2 | 49200.000 | PPDB |
| Tritosulfuron BH 635-2 | M | 1869-24-5 | 100000.000 | PPDB |
| Flurprimidol | P | 56425-91-3 | 11024.518 | EPA ECOTOX |
| Trinexapac-ethyl | P | 95266-40-3 | 142500.000 | PPDB |

**Supplementary Table 4: Contributions to $TU_{sum}$ by pesticide type and insecticide class across monitoring campaigns.** Values show the mean contribution across imputations and the corresponding 2.5th–97.5th percentile range. The final column gives the mean contribution attributable to imputed concentrations.

| Campaign | Type | class | Mean (%) | $2.5^{th}$ percentile (%) | $97.5^{th}$ percentile (%) | Mean (imputed) (%) |
|---|---|---|---|---|---|---|
| 1998–2000 | F | | 4 | 1 | 8 | 1 |
| 1998–2000 | H | | 6 | 2 | 11 | 1 |
| 1998–2000 | I | Organophosphate | 34 | 17 | 50 | 17 |
| 1998–2000 | I | Carbamate | 5 | 1 | 10 | 3 |
| 1998–2000 | I | Pyrethroid | 51 | 35 | 66 | 51 |
| 1998–2000 | I | Other | 0 | 0 | 2 | 0 |
| 1998–2000 | M | | 0 | 0 | 0 | 0 |
| 2013 | F | | 10 | 6 | 14 | 1 |
| 2013 | H | | 2 | 1 | 4 | 0 |
| 2013 | I | Organophosphate | 2 | 0 | 7 | 2 |

| | | | | | | |
|---|---|---|---|---|---|---|
| 2013 | I | Carbamate | 1 | 0 | 5 | 1 |
| 2013 | I | Pyrethroid | 19 | 12 | 29 | 10 |
| 2013 | I | Neonicotinoid | 66 | 58 | 74 | 15 |
| 2013 | M | | 0 | 0 | 1 | 0 |
| 2018–2019 | F | | 3 | 3 | 3 | 0 |
| 2018–2019 | H | | 3 | 3 | 3 | 0 |
| 2018–2019 | I | Organophosphate | 0 | 0 | 0 | 0 |
| 2018–2019 | I | Carbamate | 7 | 7 | 7 | 0 |
| 2018–2019 | I | Neonicotinoid | 72 | 71 | 72 | 0 |
| 2018–2019 | I | Other | 13 | 13 | 13 | 0 |
| 2018–2019 | M | | 2 | 2 | 2 | 0 |
| 2021 | F | | 1 | 1 | 2 | 0 |
| 2021 | H | | 2 | 1 | 2 | 0 |
| 2021 | I | Organophosphate | 0 | 0 | 0 | 0 |
| 2021 | I | Carbamate | 13 | 12 | 13 | 0 |
| 2021 | I | Pyrethroid | 50 | 48 | 53 | 3 |
| 2021 | I | Neonicotinoid | 22 | 20 | 24 | 2 |
| 2021 | I | Other | 12 | 12 | 13 | 1 |
| 2021 | M | | 0 | 0 | 0 | 0 |

**Supplementary Table 5: Contributions to $TU_{sum}$ by regulatory status across monitoring campaigns.** Values show the mean contribution across imputations and the corresponding 2.5th–97.5th percentile range. The final column gives the mean contribution attributable to imputed concentrations.

| Campaign | Status | Mean (%) | $2.5^{th}$ percentile (%) | $97.5^{th}$ percentile (%) | Mean (imputed) (%) |
|---|---|---|---|---|---|
| 1998–2000 | authorised | 100 | 100 | 100 | 74 |
| 2013 | authorised | 79 | 70 | 87 | 19 |
| 2013 | authorised (extension) | 17 | 12 | 24 | 7 |
| 2013 | not authorised | 4 | 0 | 13 | 3 |

| | | | | | |
|---|---|---|---|---|---|
| 2018–2019 | authorised | 7 | 7 | 7 | 0 |
| 2018–2019 | authorised (extension) | 66 | 65 | 66 | 0 |
| 2018–2019 | not authorised | 25 | 24 | 25 | 0 |
| 2021 | authorised | 46 | 41 | 49 | 3 |
| 2021 | authorised (extension) | 29 | 27 | 32 | 2 |
| 2021 | not authorised | 25 | 22 | 28 | 2 |
